# Immune Infiltration and Survival Analysis in Lung Adenocarcinoma: Identifying Prognostic Cell Types

**DOI:** 10.1101/2024.05.11.593151

**Authors:** Madhulika Verma

**Affiliations:** Jawaharlal Nehru University

## Abstract

The tumor microenvironment (TME) plays a crucial role in the development and survival of neoplastic cells, with tumor-infiltrating leukocytes (TILs) constituting a significant component. This immune infiltrate exhibits a diverse composition of adaptive and innate immunological cell subtypes, with varying prognostic implications across different cancer types. Recent advancements in immunotherapy underscore the importance of evaluating TILs as potential biological identifiers, particularly in the context of novel treatment strategies. In lung adenocarcinoma, the most prevalent histological subtype of lung cancer, multiple immune cell types have been identified within the TME, influencing tumor classification, clinical outcomes, and patient survival. While prior research has demonstrated a correlation between tumor-infiltrating immune cells and the progression of lung adenocarcinoma, few studies have examined their prognostic implications comprehensively.

Building upon our previous work, where we constructed a signature matrix (Verma, 2024b.) and evaluated the fractions of 14 immune cell types in TCGA-LUAD data and performed ESTIMATE analysis to assess immune infiltration, stromal infiltration, and tumor purity (Verma, 2024a.), in this study, we investigate the association between immune cell infiltration patterns and the overall survival and prognosis of TCGA-LUAD patients across different histological subtypes and stages. Our findings aim to elucidate the immune cell types positively or negatively impacting patient outcomes in lung adenocarcinoma and inform future therapeutic approaches.

## 1. Introduction

The tumor microenvironment (TME) is a complex milieu essential for the development and sustenance of neoplastic cells. Comprising diverse non-tumor components, including tumor-infiltrating leukocytes (TILs), the TME orchestrates intricate interactions that influence tumor progression and therapeutic responses. TILs represent a heterogeneous population of immune cells, encompassing various adaptive and innate immunological subtypes with distinct functionalities. While cytotoxic T cells exhibit proactive antitumor effects, regulatory T cells (T-regs) and myeloid-derived suppressor cells exert immunosuppressive functions, shaping the immunological landscape within the TME (Fridman et al., 2012; Gentles et al., 2015).

The significance of TILs in cancer pathogenesis varies across different malignancies, with specific immune populations demonstrating divergent prognostic implications. Recent advancements in immunotherapy have underscored the importance of evaluating TILs as potential biomarkers for therapeutic response prediction and patient stratification. Notably, studies have highlighted the prognostic relevance of T cell subsets, such as CD8+, in prognosticating the efficacy of immunotherapeutic interventions, emphasizing the critical role of immune cell profiling in cancer management (Tumeh et al., 2014; Herbst et al., 2014; Ji et al., 2011).

Lung adenocarcinoma, the predominant histological subtype of lung cancer, exhibits intricate interactions with various immune cell populations within the TME. Studies have elucidated the infiltration of multiple immune cell types, including T cells, B lymphocytes, dendritic cells (DCs), natural killer (NK) cells, and macrophages, into the lung TME, influencing tumor classification, clinical outcomes, and patient survival (Zheng et al., 2017; Bremnes et al., 2016; Domingues et al., 2016). Notably, the composition and proportion of these immune cells infiltrating tumors significantly impact the surrounding tissue milieu, underscoring their role as key discriminators of tumor classification and prognosticators of patient outcomes. While prior research has established a correlation between tumor-infiltrating immune cells and lung adenocarcinoma progression, limited studies have comprehensively examined their prognostic implications. We have endeavored to address this gap in the present paper.

In our prior investigation, we assessed the proportions of 14 distinct immune cell types within TCGA-LUAD datasets. Additionally, ESTIMATE analysis was conducted to quantify the extent of immune cell infiltration, stromal infiltration, and tumor purity (Verma, 2024a.).

In this study, we aim to elucidate the association between these immune cell profiles and the overall survival of TCGA-LUAD patients. Specifically, we endeavor to identify the immune cell types that positively or negatively correlate with the prognosis and survival outcomes of patients across various histological subtypes and disease stages.

The signature matrix used to deconvolute the TCGA-LUAD data is provided as supplementary file in the first paper (Verma, 2024b.) and the results of the deconvolution and the ESTIMATE analysis of TCGA-LUAD data are provided as the supplementary files in the second paper (Verma, 2024a.).

## 2. Materials and Methods

### 2.1. Data

The “TCGAbiolinks” (Colaprico et al., 2016) R package was used to get lung adenocarcinoma datasets from the TCGA (*The Cancer Genome Atlas Program*, n.d.). The category of the data was “Transcriptome Profiling”, the data_type was “Gene Expression Quantification”, and the workflow_type was “STAR - Counts”. These were the technical terms that were used to extract the desired data. The data consisted of 60,660 genes from 598 patients (samples). The other specifications for this data were as follows:

- Out of 598 samples:

○ Primary Tumor - 537
○ Recurrent Tumor - 2
○ Solid Tissue Normal - 59
- Out of 598, the vital status:

○ Alive - 380
○ Dead - 218
- Out of 598, pathologic_stage

○ Stage I - 5
○ Stage IA - 152
○ Stage IB - 169
○ Stage II - 1
○ Stage IIA - 55
○ Stage IIB - 82
○ Stage IIIA - 85
○ Stage IIIB - 12
○ Stage IV - 28
- Primary Diagnosis (Histological subtypes)

○ Acinar cell carcinoma - 22
○ Adenocarcinoma with mixed subtypes - 114
○ Adenocarcinoma, NOS - 374
○ Bronchio-alveolar carcinoma, mucinous - 5
○ Bronchiolo-alveolar adenocarcinoma, NOS – 3
○ Bronchiolo-alveolar carcinoma, non-mucinous - 22
○ Clear cell adenocarcinoma, NOS - 3
○ Micropapillary carcinoma, NOS - 3
○ Mucinous adenocarcinoma - 20
○ Papillary adenocarcinoma, NOS - 25
○ Signet ring cell carcinoma - 1
○ Solid carcinoma, NOS - 6

In addition to these details, the data had full clinical details, such as whether or not the person smoked, the molecular subtype, the type of treatment, the person’s age, gender, race, and many more.

### 2.2. Extracting the clinical data

The clinical data that went with the TCGA-LUAD data included information like patient barcodes, stages, histological subtypes, treatments, smoking status, date of diagnosis, status as alive or dead, weight, age, gender, and a lot of other information that was clinically important. For our analysis, we used the following information:

**Table 1.** Clinical variables and their meaning (description as per the TCGA data dictionary)

| Variables | Meaning |
| --- | --- |
| patient | Patient ID in the study |
| sample | Unique sample barcode of patients |
| age_at_diagnosis | “Age at the time of diagnosis expressed in number of days since birth.” |
| ajcc_pathologic_stage | “The extent of cancer, especially whether the disease has spread from the original site to other parts of the body based on AJCC staging criteria.” |
| days_to_diagnosis | “Number of days between the date used for index and the date the patient was diagnosed with the malignant disease.” |
| days_to_last_follow_up | “Time interval from the date of last follow-up to the date of initial pathologic diagnosis, represented as a calculated number of days.” |
| primary_diagnosis | “Text term used to describe the patient's histologic diagnosis, as described by the World Health Organization's (WHO) International Classification of Diseases for Oncology (ICD-O).” |
| gender | “Gender of the patient” |
| year_of_diagnosis | “Numeric value to represent the year of an individual's initial pathologic diagnosis of cancer.” |
| year_of_death | “Numeric value to represent the year of the death of an individual”. |
| days_to_death | “Number of days between the date used for index and the date from a person's date of death represented as a calculated number of days.” |
| vital_status | “The survival state of the person registered on the protocol.” |

Apart from the above information, we also added the scores of the 14 immune cell types as columns to the above clinical data, along with the ESTIMATE score, immune score, and stromal score. This clinical data was organized so that the rows represented the samples and the columns represented the above-mentioned clinical information and cell type scores for each sample.

### 2.3. Classification of the data (histotypes and stages)

After retrieving the clinical data, the tumor samples were sorted. In the TCGA-LUAD data, out of a total of 598 samples, 537 were from the primary tumor, 2 were from the recurrent tumor, and 59 were adjacent normals. Since there were only 2 recurrent samples, we could not perform the disease-free survival analysis with such a small number; therefore, we only concentrated on the primary tumor, which had a sufficiently large number of samples for calculating overall survival. We separated the primary tumor sample and the corresponding clinical data to calculate the overall survival.

The primary tumor samples were divided into 12 different histological subtypes listed under the heading “primary_diagnosis” in the clinical data, and 4 stages (and their substages). We were interested in finding out the effect of the individual immune cell level, the total immune and stromal content, and the tumor purity on the primary tumor data as well as its constituent subtypes and stages. For this purpose, we divided the primary tumor samples into 12 histotypes along with the corresponding clinical data. Out of these 12 histo-types, the Acinar cell carcinoma, Adenocarcinoma with mixed subtypes, Adenocarcinom-NOS, Bronchiolo-alveolar carcinoma (non-mucinous), Mucinous adenocarcinoma, Papillary adenocarcinoma-NOS, had a sufficient number of samples for a reliable survival analysis. The remaining subtypes Bronchio-alveolar carcinoma-mucinous (5), Bronchiolo-alveolar adenocarcinoma-NOS (3), Clear cell adenocarcinoma-NOS (3), Micropapillary carcinoma-NOS (3) Signet ring cell carcinoma (1), and Solid carcinoma-NOS (6), had a very small number of samples thus, all these samples were merged together to form a group named “mixed.” This mixed group had 21 samples, to be precise. Similarly, the primary tumor was also divided into four stages. The clinical data also showed that there were substages, so the substages were added to the main stages they belonged to. For example, Stages I, IA, and IB were merged together to form Stage I; Stages II, IIA, and IIB were combined to form Stage II; Stages III, IIIA, and IIIB were combined to form Stage III; and Stage IV was the final stage. Thus, we got our stages from the primary tumor. The clinical data were also separated accordingly for all histotypes and stages.

### 2.4. Survival analysis and plots (histotypes and stages)

Once we got our data in place, we went on with the actual survival estimate. The packages “survival” (v3.4-0, Therneau, 2022, Borgan, 2001) and “survminer” (Kassambara et al., 2021) from R were used for this purpose. The important variables needed for the survival analysis are the patient’s status--whether or not the patient is alive, pathologic stage (4 stages), and 2 crucial pieces of information: days_to_death, which is the count, in days, from the time of diagnosis till the patient passed away, and days_to_last_follow-up, which is the time period between the original diagnosis and the patient’s final consultation. We changed some of this information so that it would work with the procedures in the package “survival”. We generated a new binary instance with FALSE for those who were still alive and TRUE for deceased subjects. Then, we constructed the “overall survival” instance, which consisted of days_to_death for deceased subjects and days_to_last_follow_up for living patients. Next, we added the censoring information to the patients, which means just adding the + just after the time, for all the patients. For some instances, we just have the time when they last followed up and don’t know whether they are alive or dead after that, making this data right censored. Although these patients are included in the initial phases of analysis (such as the survival curve), they are subsequently excluded (or “censored”) when the end of their follow-up period approaches. The patients that are alive and still in the study are said to have no event (death or recurrence).

### 2.5. Kaplan Meier Analysis and plots

The Kaplan-Meier (KM) method (Kaplan and Meier, 1958) is a non-assumptive way to estimate the chance of survival based on how long people have lived. We performed univariate survival analysis on categorical data using the Kaplan-Meier method. All our variables (ESTIMATE results, CIBERSORTx (Steen et al., 2020) score) were numerical data and could not be converted into factors. Therefore, first we categorized our data with numerical values into groups. For each type of score, be it from CIBERSORTx (Steen et al., 2020) for immune cells or the ESTIMATE score, we calculated the median of values and categorized the data as a high or low group depending on whether their score was higher than the median value or lower. As a result, we had high and low groups for 14 immune cell types, as well as the three ESTIMATE (Yoshihara et al., 2013) scores. This process was done for the primary tumor samples as well as the 7 histotypes and 4 stages. So now we have an extra column in our clinical data that indicates if a patient is in a high or low group for each of the above-mentioned variables. Then we defined a survival formula that took the censoring data (overall survival plus deceased information), followed by the categorical data (cell types score or stromal/immune or estimate score). The *survfit* function was then used to fit this survival formula to a survival model, and survminer was used to generate the Kaplan-Meier plots. The plots showed two curves, one for each category, showing differences in the survival probabilities of the groups, and the significance of this difference was determined by using the log-rank method, that is essentially a repeating test of no dependence. This p-value was depicted on the KM plots generated. A risk table was also generated and depicted on the KM plots; it showed the number of patients “at risk”, meaning they were not censored or dead at a certain time point.

## 3. Results and Discussion

### 3.1. Data

We had three sets of data to perform the survival analysis on. First were the primary tumor data, then the 7 histological subtypes, and finally the 4 stages. In these three datasets, we checked the effect of the CIBERSORT immune cells score, the score of purity of tumor, the immune score, and the stromal score on the overall survival of the patients.

### 3.2. Primary Tumor

In the primary tumor, three cell types—activated Activated/matured B cells (pval = 0.0043), monocyte-derived DC (pval = 0.034), and naive CD4 T cells (pval = 0.007)—had a substantial impact on the overall survival of lung adenocarcinoma patients. Patients with a high level of activated or mature B cells have a better chance of survival than those with a low level. Similarly, the patients with high levels of CD4 T cells had a higher overall survival probability than the patients with low levels. The same thing was observed with monocyte-derived DCs: cases with greater DC content had a higher probability of survival than the ones with lower levels.

When the ESTIMATE scores were observed in the primary tumor, although patients with lower immune scores seemed to have greater risk over time, the model was insignificant. The result was inconclusive (pval = 0.86).

**Figure 1.**
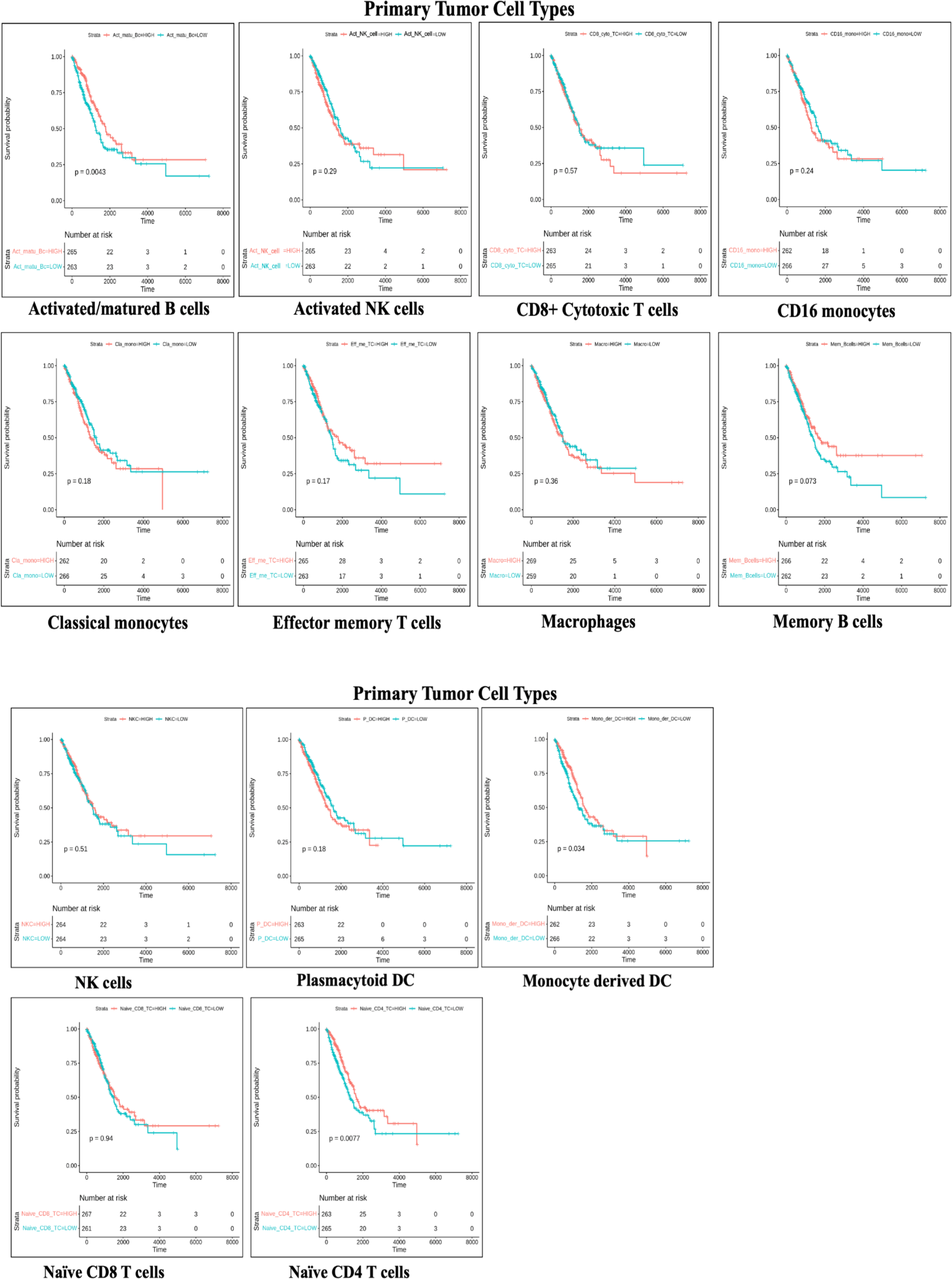
Kaplan-Meier plots showing the effect of 14 immune cell types on the overall survival of LUAD patients with primary tumors (5 out of 14). Three cell types (Activated/matured B cells, monocyte-derived DC, and naive CD4 T cells) have a significant effect on the overall survival of LUAD patients.

**Figure 2.**
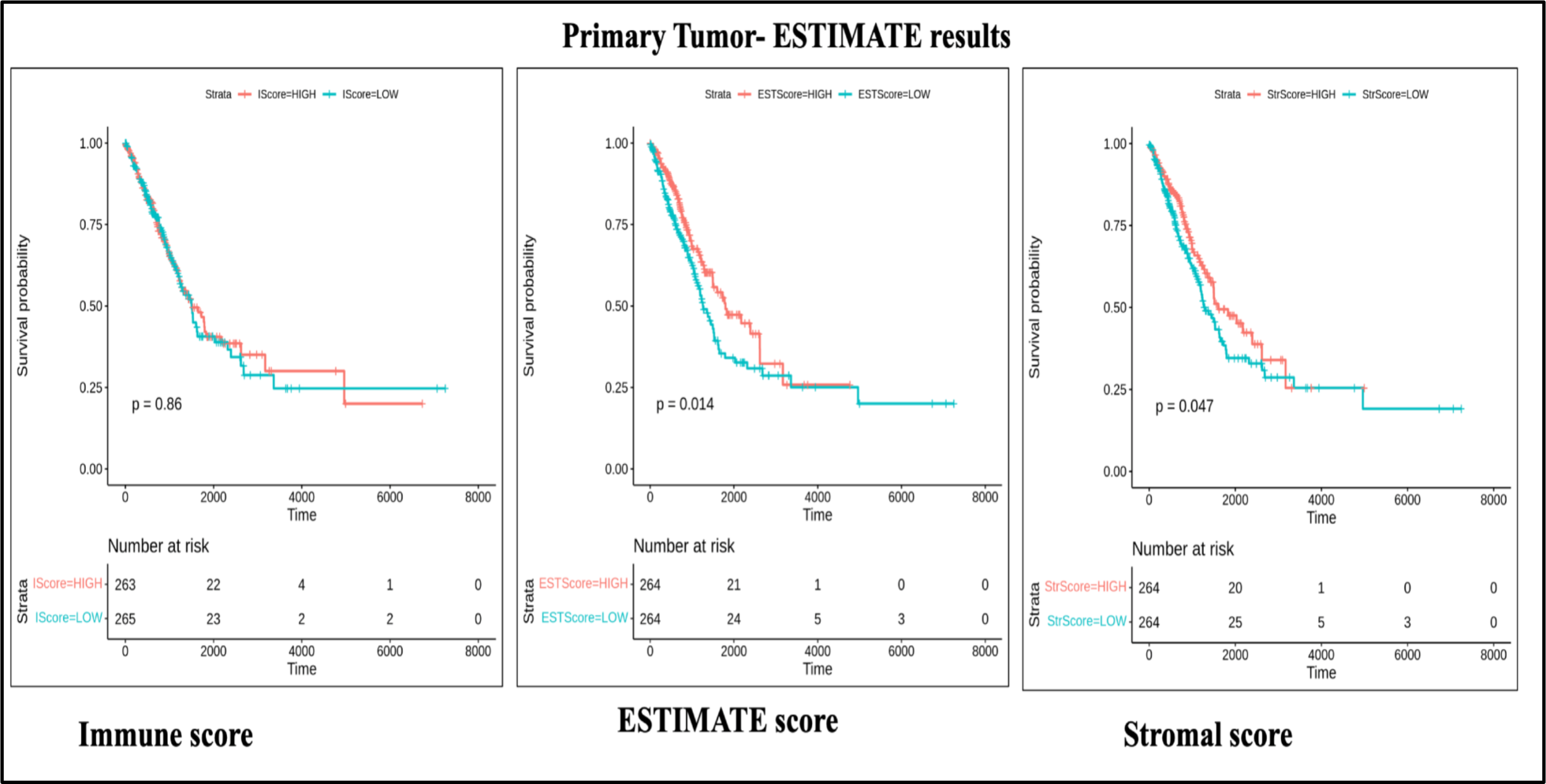
Kaplan-Meier curves showing the effect of the ESTIMATE-derived scores on the overall survival of the LUAD patients with primary tumors. Stromal and ESTIMATE scores show a significant difference in overall survival between the low-score and high-score groups.

Stromal score showed a significant (pval = 0.047) difference in survival time between the group having a lower stromal score and the group having a higher stromal score along with the higher and lower ESTIMATE score (tumor purity) groups had same kinds of results (pval = 0.014) as the high and low stromal score groups.

### 3.3. Histological Subtypes

The data from TCGA-LUAD showed that there were 12 histological subtypes. By combining these subtypes, as explained in the “Methods” section, we were able to get 7 major histological groups. The first was the acinar cell carcinoma subtype, in which only the NK cells (pval= 0.012) showed a substantial variation in the overall survival among the low and high category of patients. None of the immune cells appeared to have a significant effect on patient survival in adenocarcinomas with mixed subtypes. High and low levels of Activated/matured B cells (pval = 0.0007), monocyte-derived DC (pval = 0.033), and plasmacytoid DC (0.016), showed significance in the overall survival of patients with Adenocarcinoma. Surprisingly, while activated/mature B cells and monocyte-derived DC increased the survival probability in the high-level group, plasmacytoid DC increased the survival probabilities in the low-level group. In the bronchiolo-alveolar-carcinoma non-mucinous subtype, only patients with high and low memory B cell subsets had a significant difference according to the statistics (pval = 0.019) in overall survival. Then again, none of the immune cells showed a noteworthy and statistically accepted difference in the overall survival of mucinous adenocarcinoma patients. In papillary adenocarcinoma subtype patients, naive CD8 T cells appeared to show a significant difference (pval = 0.003), i.e., patients with low levels of naive CD8 T cells had a lower survival probability than those with high levels. None of the immune cells show statistically significant differences in the overall survival of mixed-subtype group patients. The plots are provided in the *Appendix (Figure 0.1-0.7)*.

**Table 2.** The outcome of the survival analysis summarized, demonstrating the effect of different immune cell type fractions in various histotypes, on the overall survival of LUAD patients.

| LUAD histological subtypes | Cell type | P. value | Relation to overall survival |
| --- | --- | --- | --- |
| Acinar_cell_carcinoma | NK cells | 0.012 | Positive |
| Adenocarcinoma_with_mixed_subtypes | NA | NA | NA |
| Adenocarcinoma_NOS | Activated/matured B cells | 0.00079 | positive |
|  | Monocyte DC | 0.033 | positive |
|  | Plasmacytoid DC | 0.016 | negative |
| Bronchiolo_alveolar_carcinoma_non_mucinous | Memory B cells | 0.019 | positive |
| Mucinous adenocarcinoma | NA | NA | NA |
| Papillary adenocarcinoma | Naive CD8 T cells | 0.003 | positive |
| Mixed | NA | NA | NA |

In the case of ESTIMATE results, the immune score had a significant impact on overall survival only in adenocarcinoma patients, with a p-value of 0.024; the cases with greater immune score possessed a greater survival probability than the ones with lower immune scores. The stromal score was found to have a significant difference in the overall survival of adenocarcinoma with mixed subtype patients with a p-value of 0.027. The ESTIMATE score (tumor purity) did not have a significant effect on the overall survival of any histological subtype. The plots are provided in the *Appendix (Figure 0.8-0.10)*.

**Table 3.**
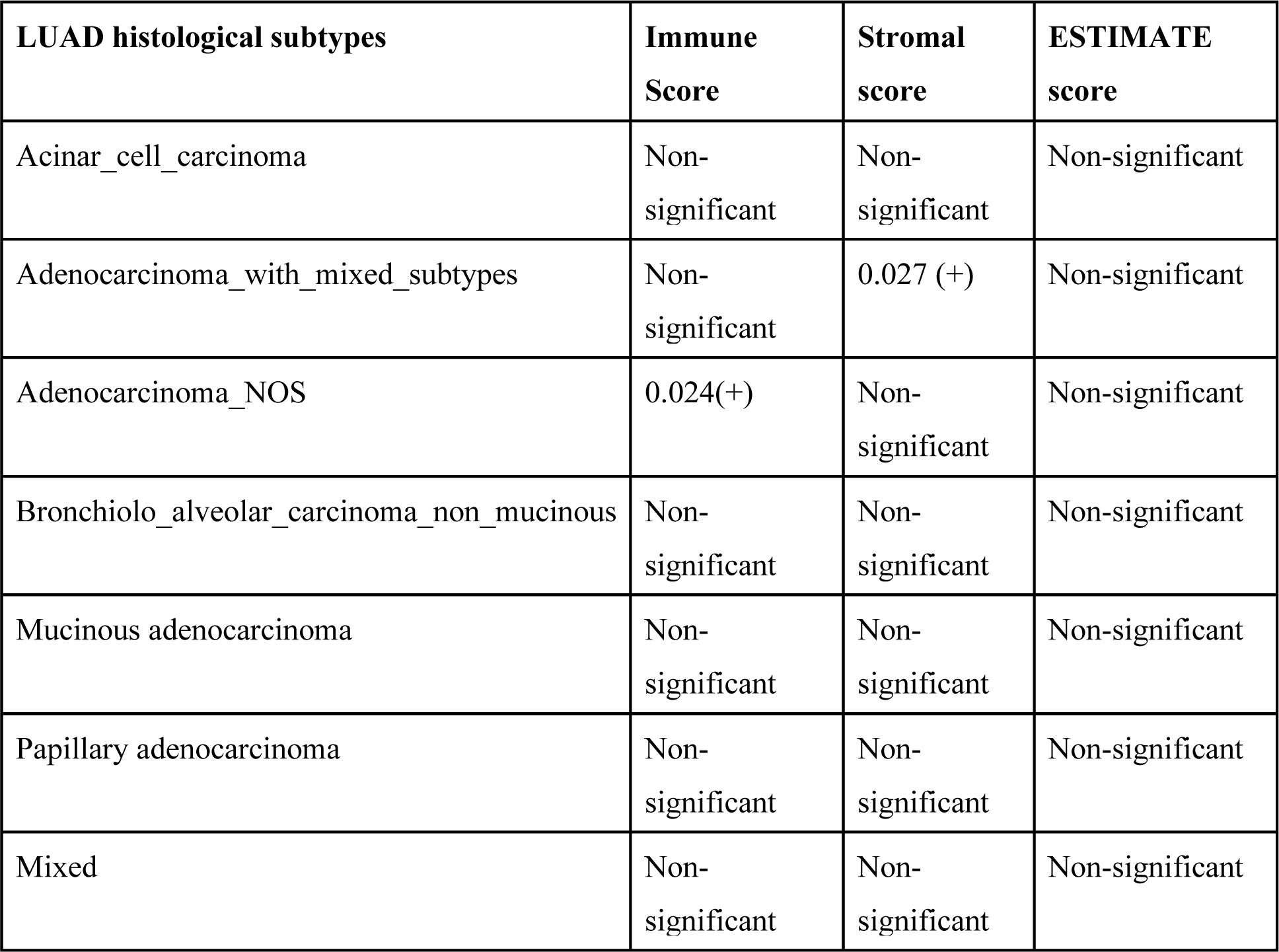
The outcome of the survival analysis summarized, demonstrating the effect of different ESTIMATE scores in various histotypes on the overall survival of LUAD patients. The numbers show the p-values, wherever the results were significant. The (+) or (-) signs depict the positive or negative correlation of the factors, with overall survival.

| **LUAD histological subtypes** | **Immune Score** | **Stromal score** | **ESTIMATE score** |
| --- | --- | --- | --- |
| Acinar_cell_carcinoma | Non-significant | Non-significant | Non-significant |
| Adenocarcinoma_with_mixed_subtypes | Non-significant | 0.027 (+) | Non-significant |
| Adenocarcinoma_NOS | 0.024(+) | Non-significant | Non-significant |
| Bronchiolo_alveolar_carcinoma_non_mucinous | Non-significant | Non-significant | Non-significant |
| Mucinous adenocarcinoma | Non-significant | Non-significant | Non-significant |
| Papillary adenocarcinoma | Non-significant | Non-significant | Non-significant |
| Mixed | Non-significant | Non-significant | Non-significant |

### 3.4. Stages

At stage I, none of the immune cells made a big difference in the patient’s overall survival, whether immune cell scores were low or high. At stage II, activated/mature B cells (pval = 0.015) and monocyte-derived DCs (pval = 0.057), had a substantial effect over the overall survival of lung adenocarcinoma subjects. In both cases, the higher the score, the better the chances of survival. At stage III, activated/mature B cells have a significant (pval = 0.042) effect on the overall survival of LUAD patients. Again, the higher the score, the greater the survival probability. Then finally, none of the immune cells showed a significant effect on the overall survival of the LUAD patients at stage IV. The plots are provided in the *Appendix (Figure 0.11-0.14)*.

**Table 4.** Summarized results of the survival analysis, demonstrating the effect of different immune cell types on the overall survival of LUAD patients at various stages of the primary tumor.

| LUAD stages | Cell types | P. value | Relation with overall survival |
| --- | --- | --- | --- |
| Stage 1 | NA | NA | NA |
| Stage 2 | Activated/matured B cells | 0.015 | positive |
|  | Monocyte DC | 0.057 | positive |
| Stage 3 | Activated/matured B cells | 0.042 | positive |
| Stage 4 | NA | NA | NA |

The immune score did not show any significance in the overall survival of the lung adenocarcinoma cases at any stage. The stromal score showed significance in the third stage. The higher level corresponded to a greater survival probability and vice versa. Only at the first stage the ESTIMATE score showed significance for overall patient survival in the greater and lower groups. Again, the higher the score, the greater the probability of survival, and vice versa The plots are provided in the *Appendix (Figure 0.15-0.17)*.

**Table 5.** Summarized results of the survival analysis, demonstrating the effect of different scores derived using ESTIMATE, on the overall survival of LUAD patients at various stages of the primary tumor. The numbers show the p-values, wherever the results were significant. The (+) or (-) signs depict the positive or negative correlation of the factors, with overall survival.

| LUAD stages | Immune Score | Stromal score | ESTIMATE score |
| --- | --- | --- | --- |
| Stage 1 | Non-significant | Non-significant | 0.056 (+) |
| Stage 2 | Non-significant | Non-significant | Non-significant |
| Stage 3 | Non-significant | 0.016 (+) | Non-significant |
| Stage 4 | Non-significant | Non-significant | Non-significant |

## 4. Conclusion

In conclusion, our study reveals several significant findings regarding the relationship between tumor-infiltrating immune cells, tumor purity, and overall survival in lung adenocarcinoma (LUAD) patients. We found that tumor purity, as indicated by the ESTIMATE score, was positively associated with overall survival, particularly in patients at the first stage of the disease. Additionally, the Stromal score showed a positive correlation with improved overall survival in patients with primary tumors, adenocarcinoma with mixed subtypes, and LUAD at stage 3. Moreover, the immune score was positively correlated with overall survival in adenocarcinoma patients.

Furthermore, our analysis identified specific immune cell types that were associated with either positive or negative overall survival outcomes. Notably, the presence of Naive CD4 T cells, Naive CD8 T cells, Memory B cells, Activated/mature B cells, NK cells, and Monocytic DCs was positively linked with overall survival in LUAD patients. Conversely, Plasmacytoid DCs were found to be negatively associated with overall survival in LUAD patients.

Overall, our findings underscore the importance of considering tumor-infiltrating immune cells and tumor purity in predicting overall survival outcomes in LUAD patients. By elucidating these associations, our study contributes to a deeper understanding of the immune landscape of LUAD and may inform the development of more effective prognostic and therapeutic strategies for this deadly disease.

## Appendix

### Survival analysis plots

**Figure 0.1.**
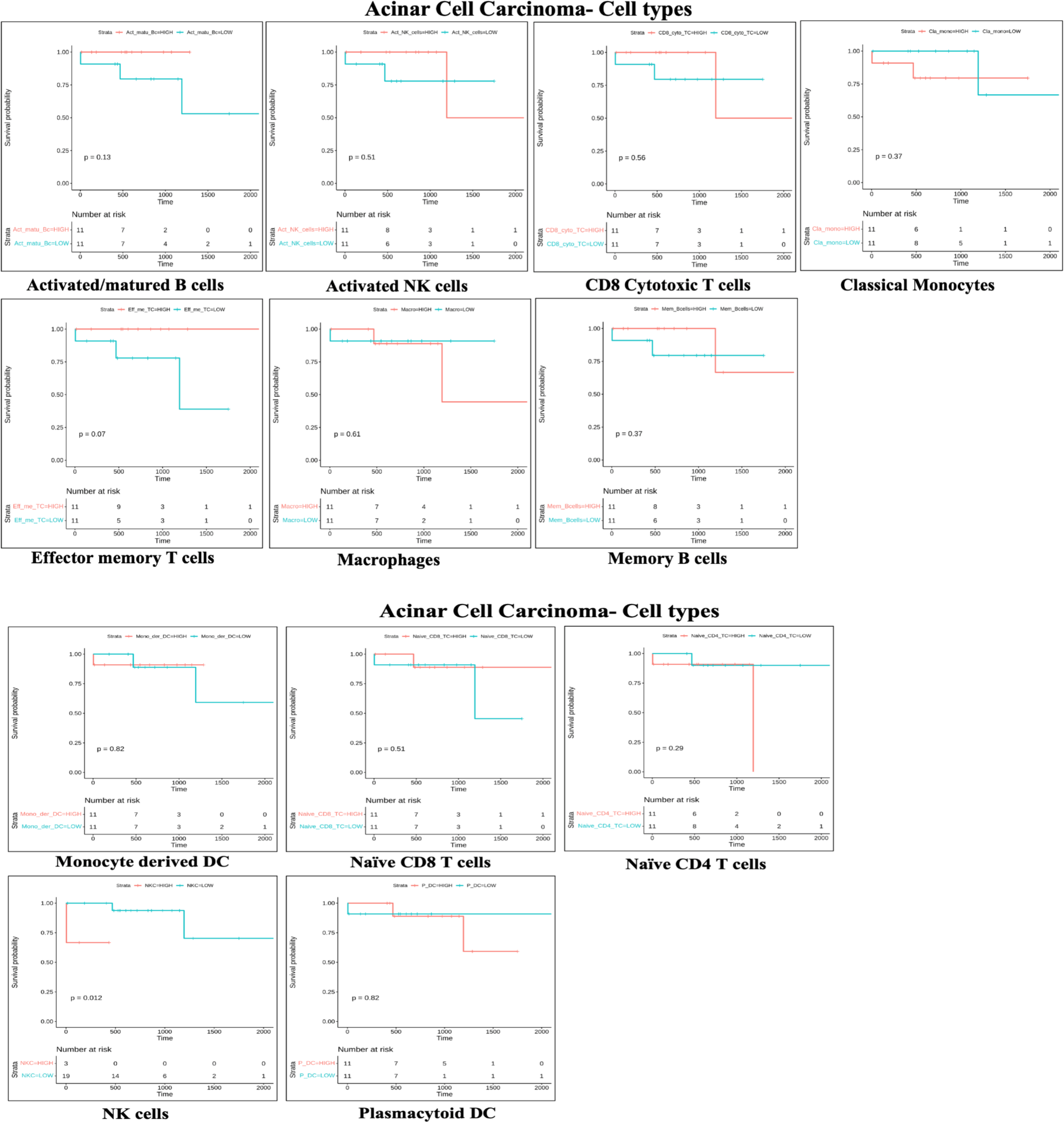
KM plot for immune cell types in Acinar cell carcinoma.

**Figure 0.2.**
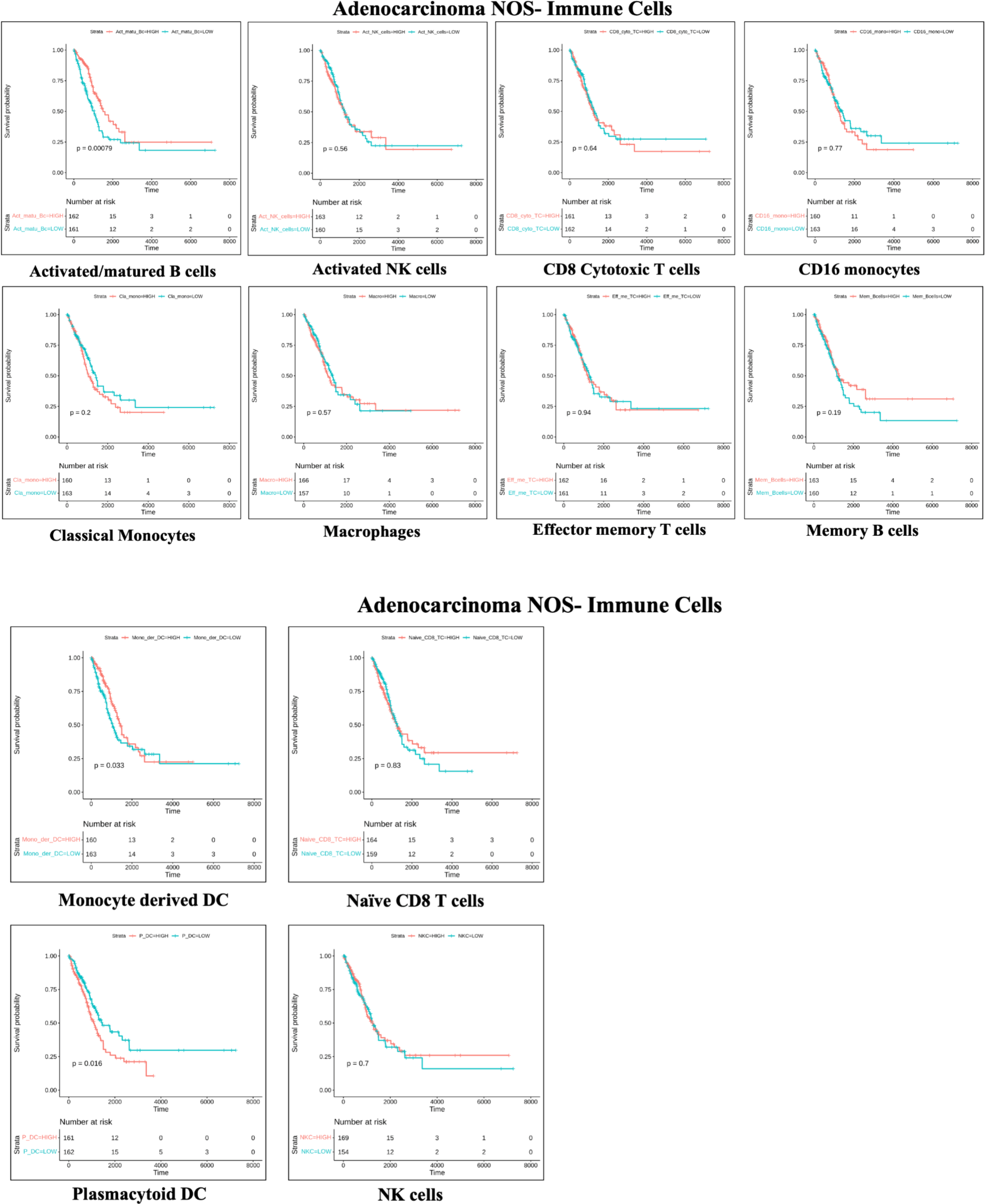
KM plot for immune cells in Adenocarcinoma NOS.

**Figure 0.3.**
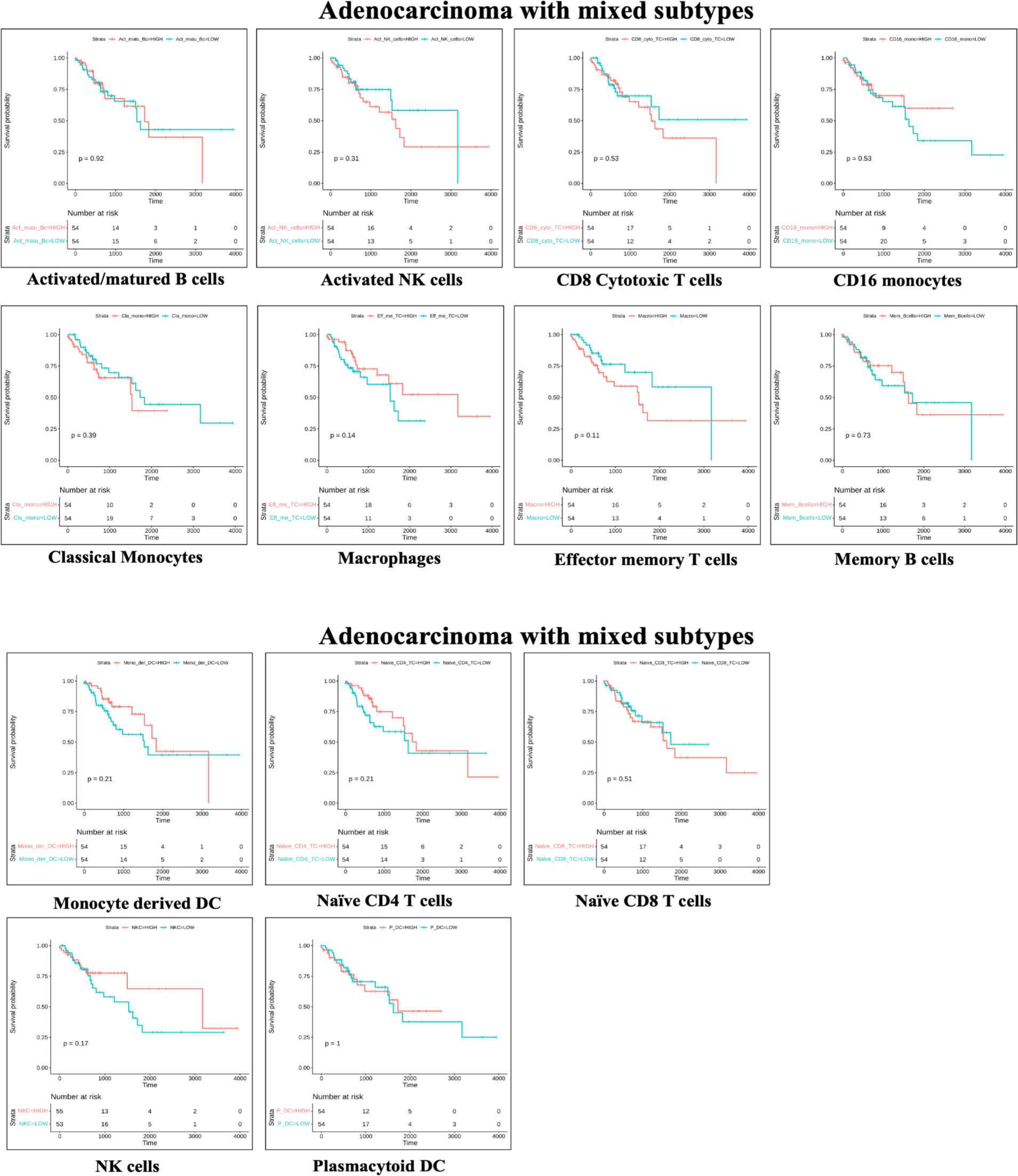
KM plot for immune cells in Adenocarcinoma with mixed subtypes.

**Figure 0.4.**
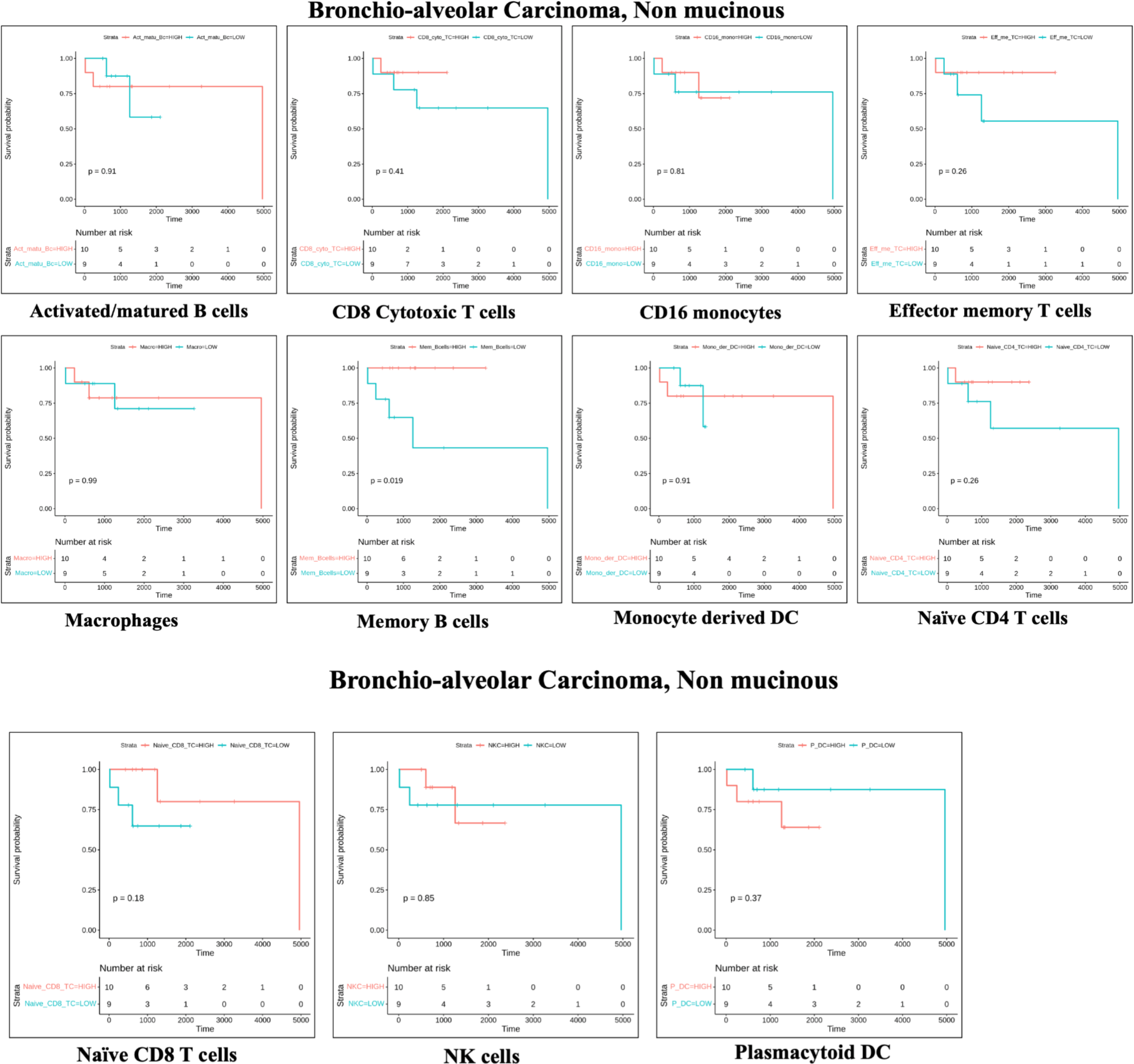
KM plot for immune cells in Bronchio-alveolar carcinoma, non-mucinous.

**Figure 0.5.**
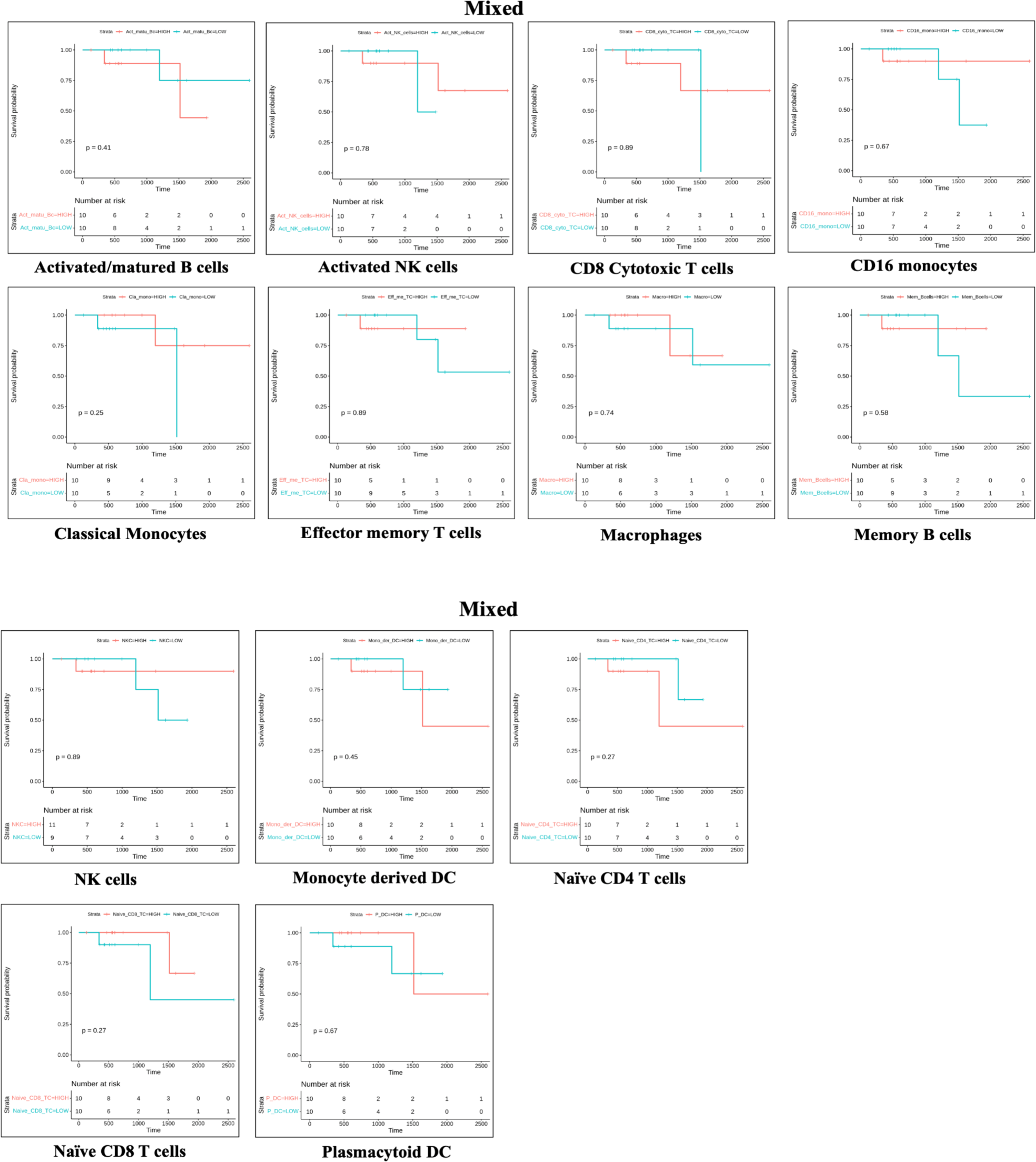
KM plot for immune cells in mixed subtypes.

**Figure 0.6.**
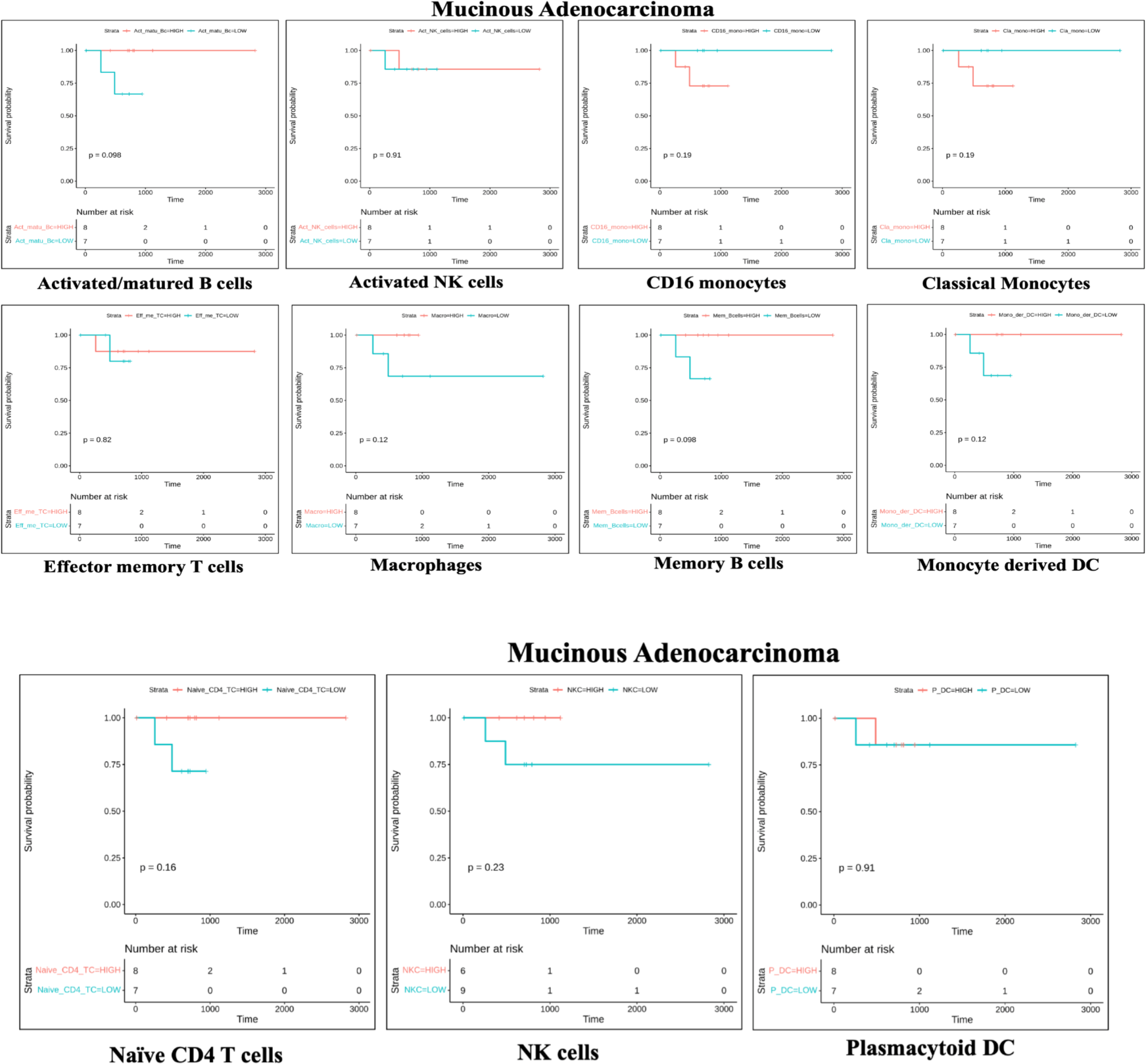
KM plot for immune cells in Mucinous adenocarcinoma.

**Figure 0.7.**
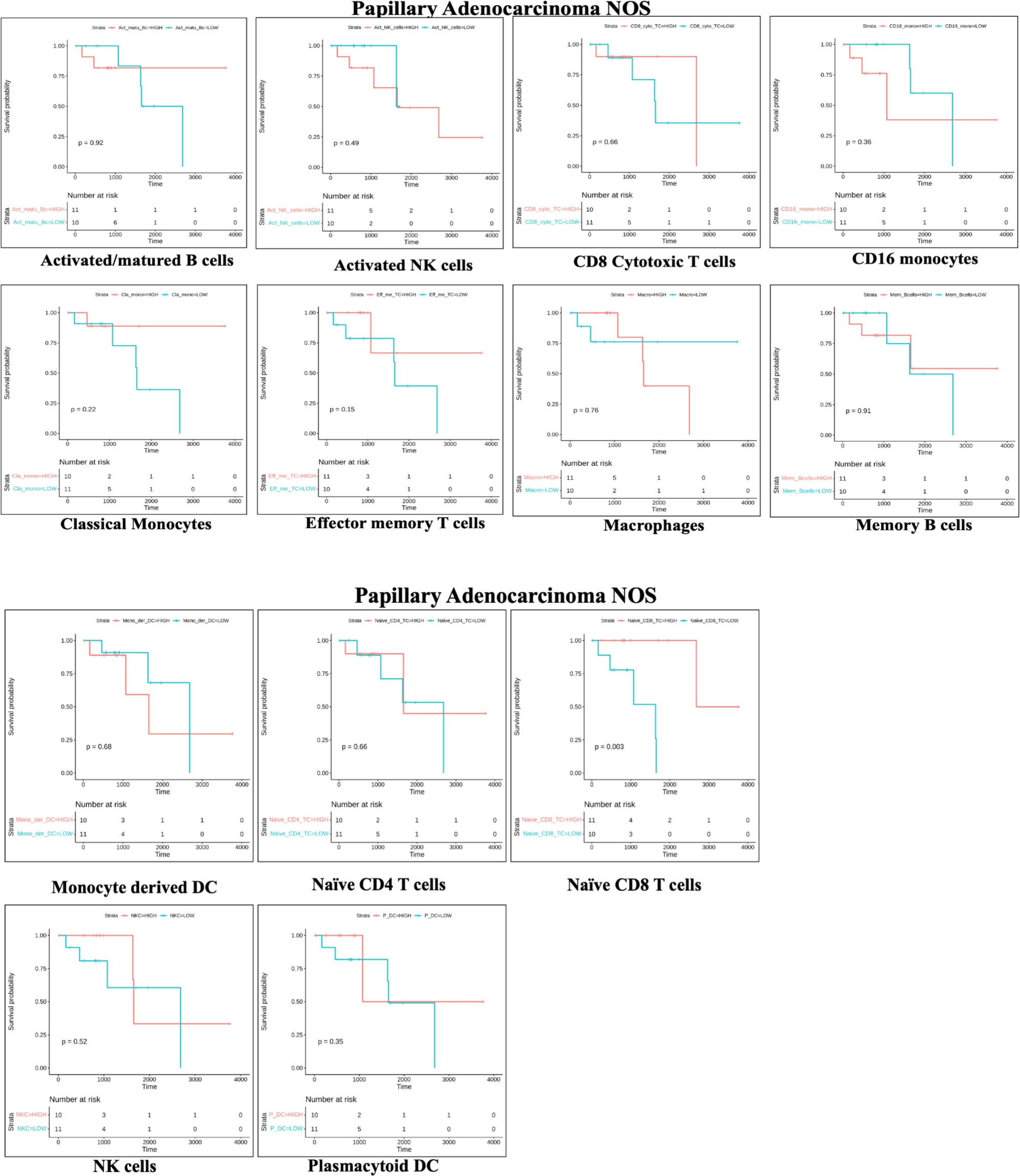
KM plot for immune cells in Papillary adenocarcinoma.

**Figure 0.8.**
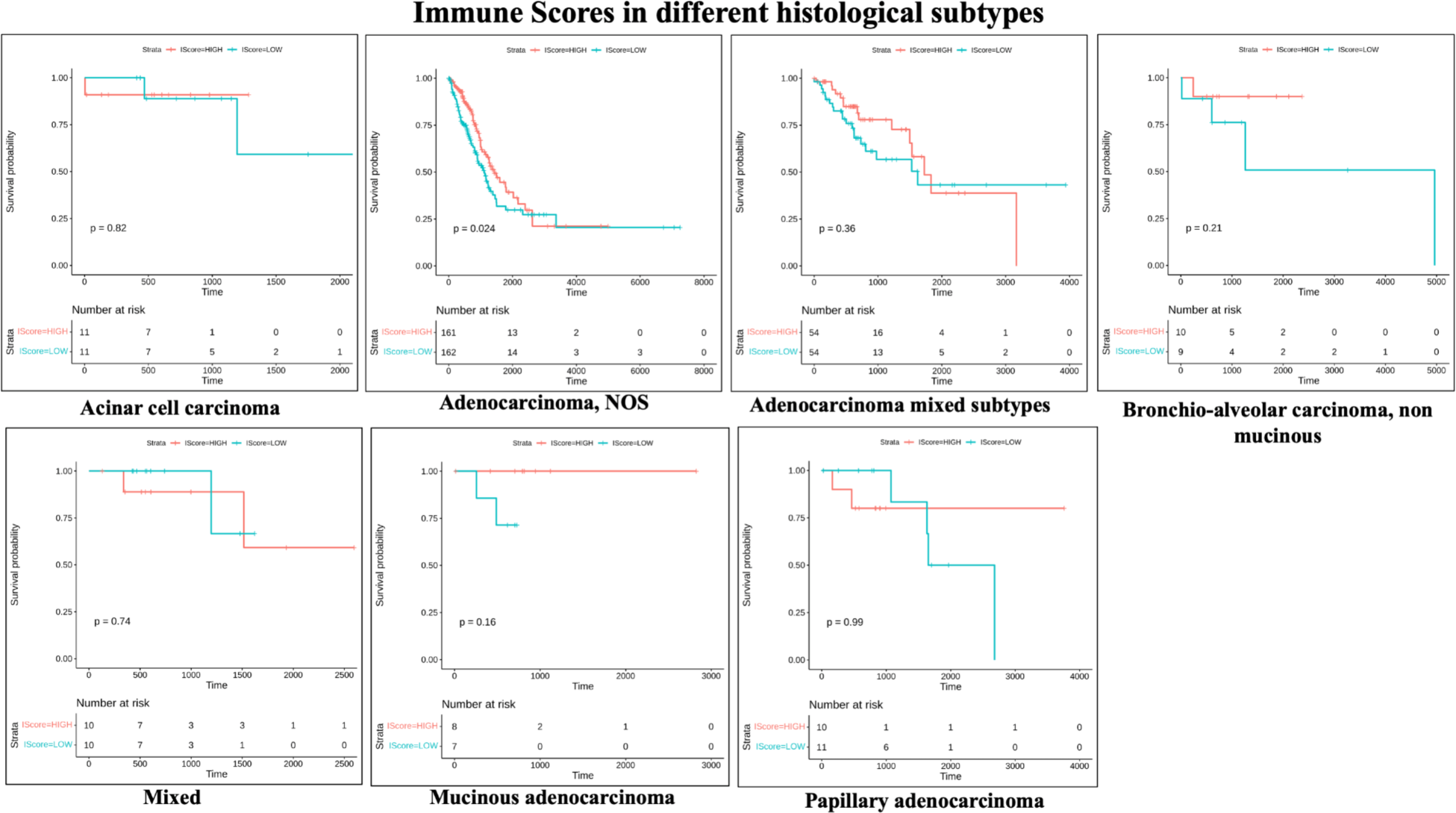
KM plot for immune scores in different histological subtypes.

**Figure 0.9.**
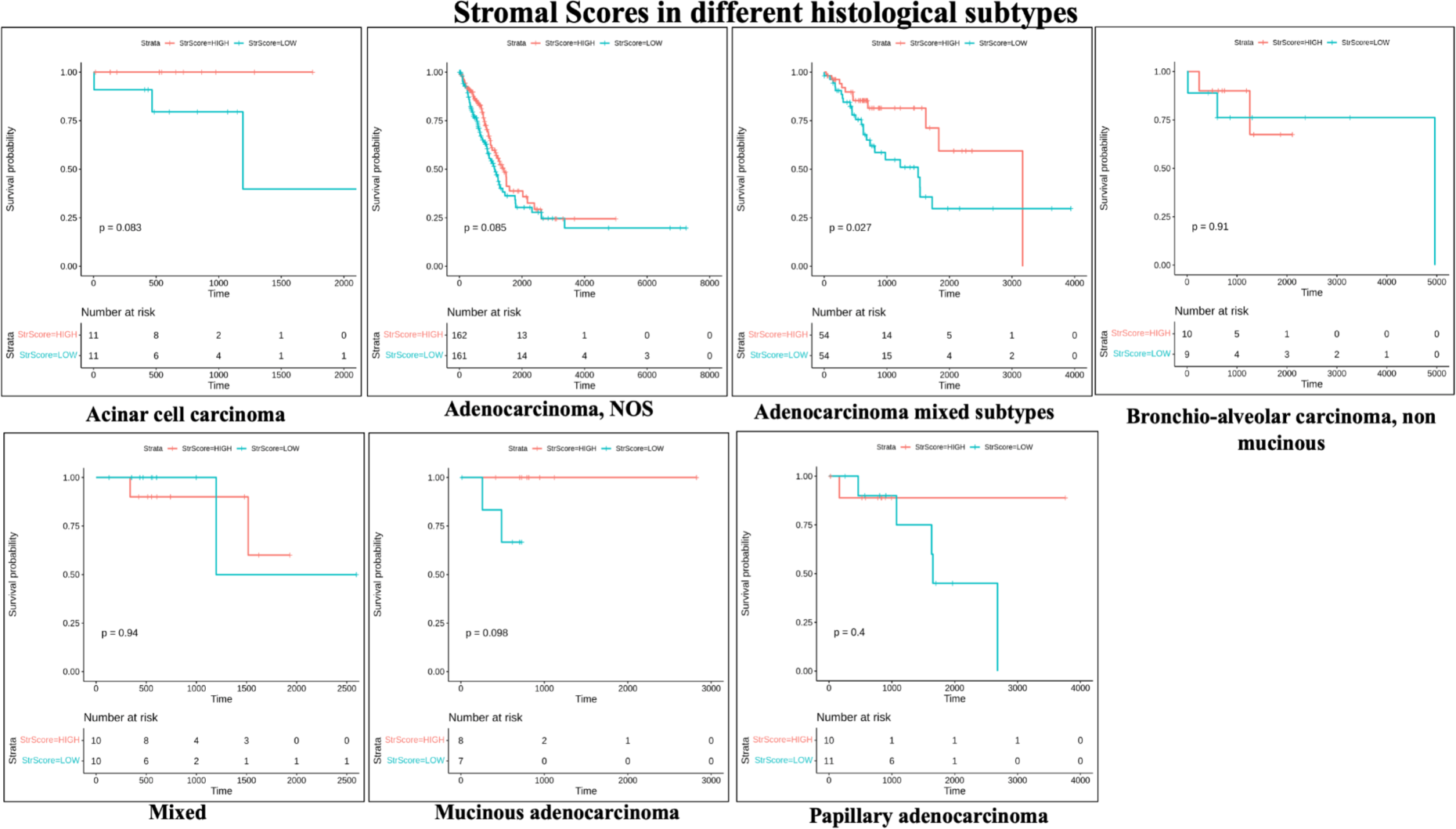
KM plot for stromal scores in different histological subtypes.

**Figure 0.10.**
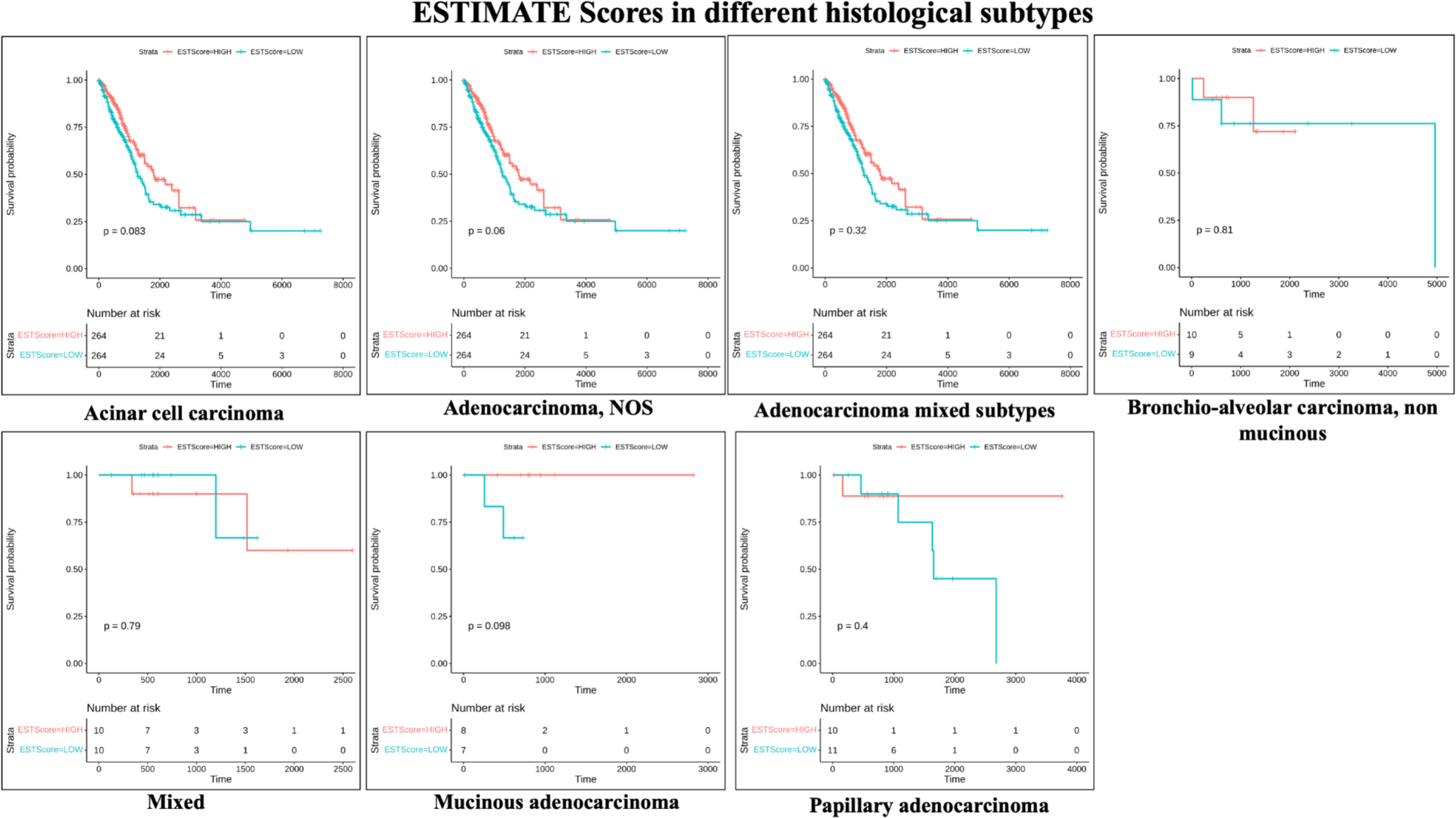
KM plot for ESTIMATE scores in different histological subtypes.

**Figure 0.11.**
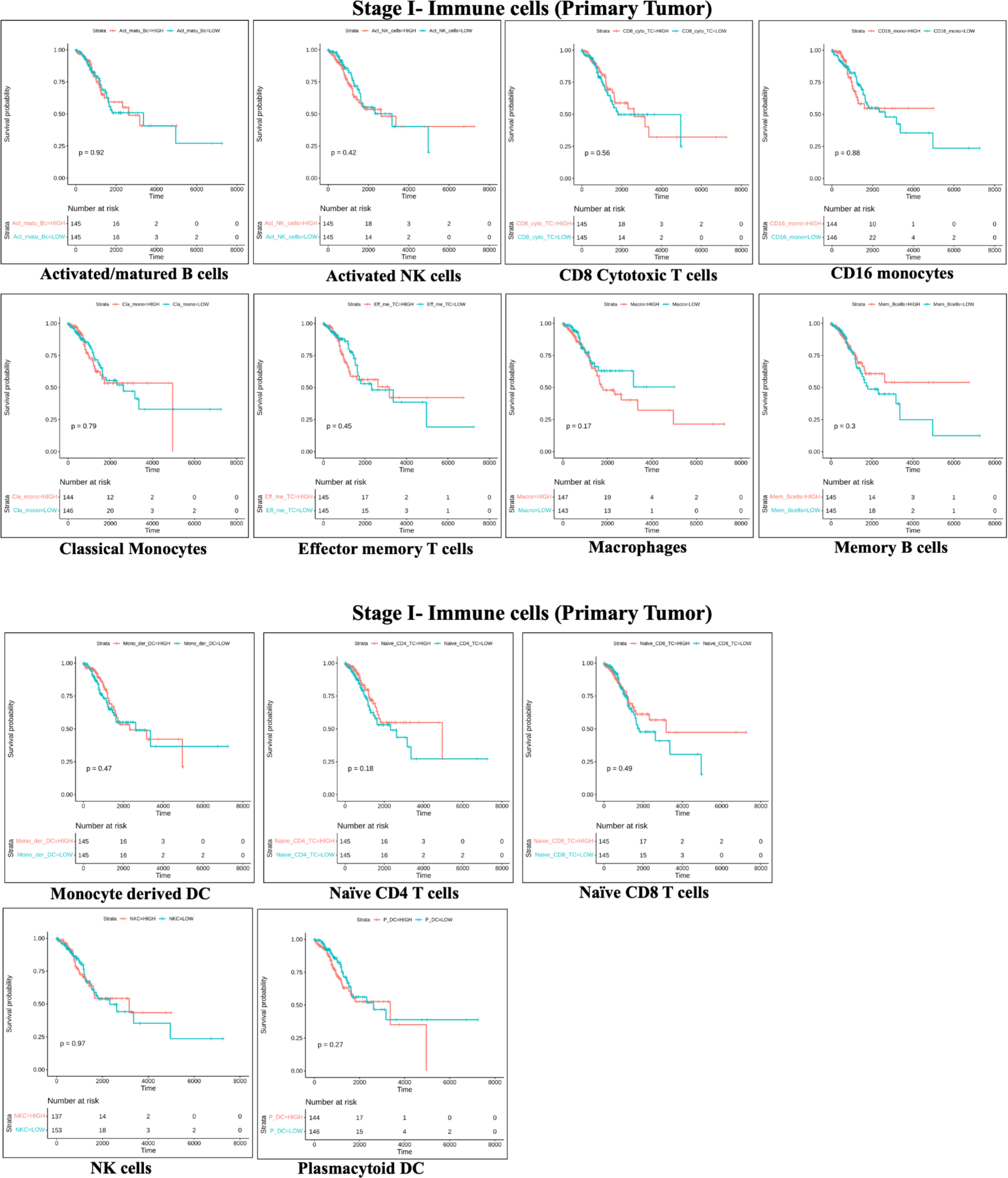
KM plot for immune cells in stage I of primary tumor.

**Figure 0.12.**
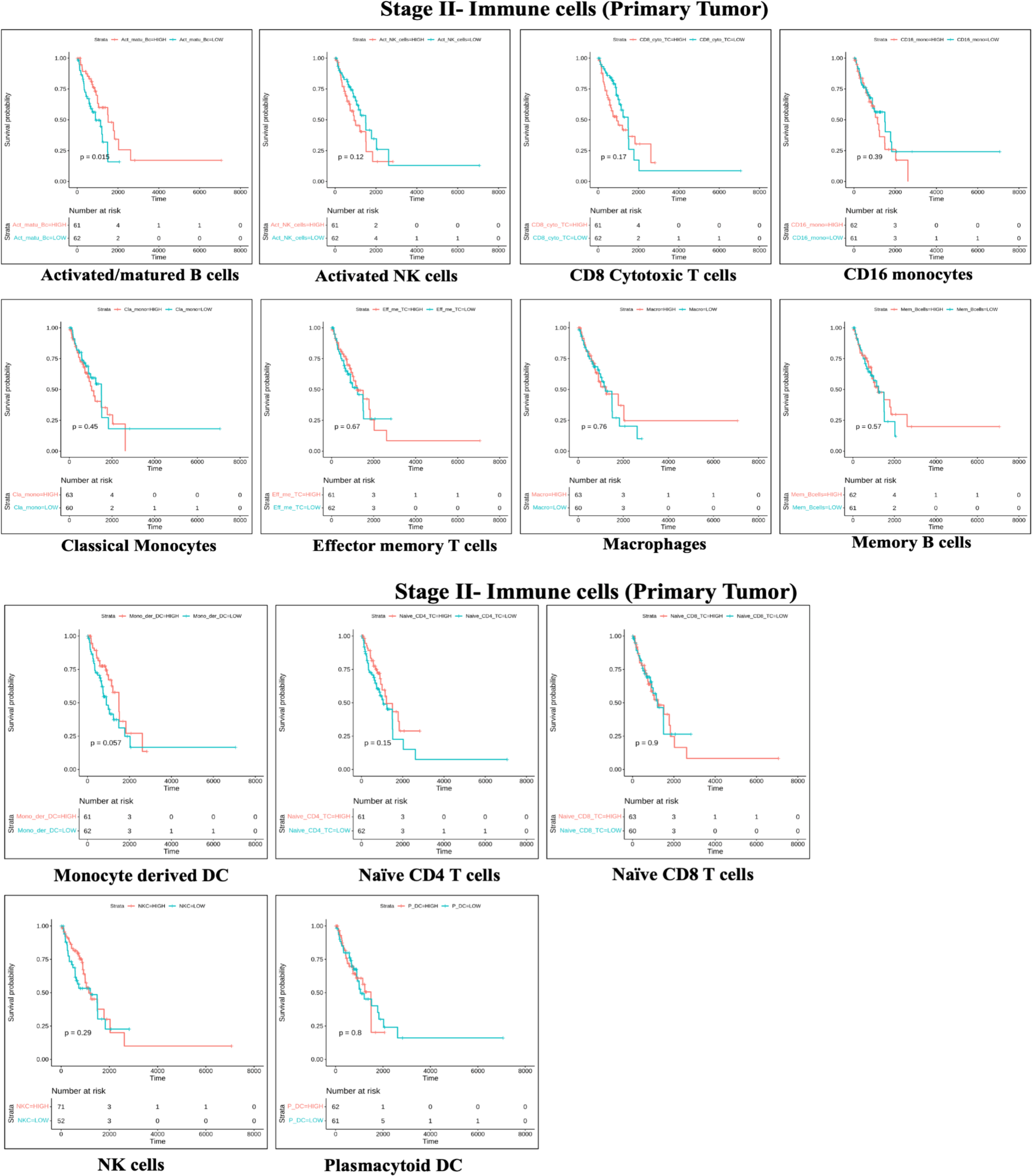
KM plot for immune cells in stage II of primary tumor.

**Figure 0.13.**
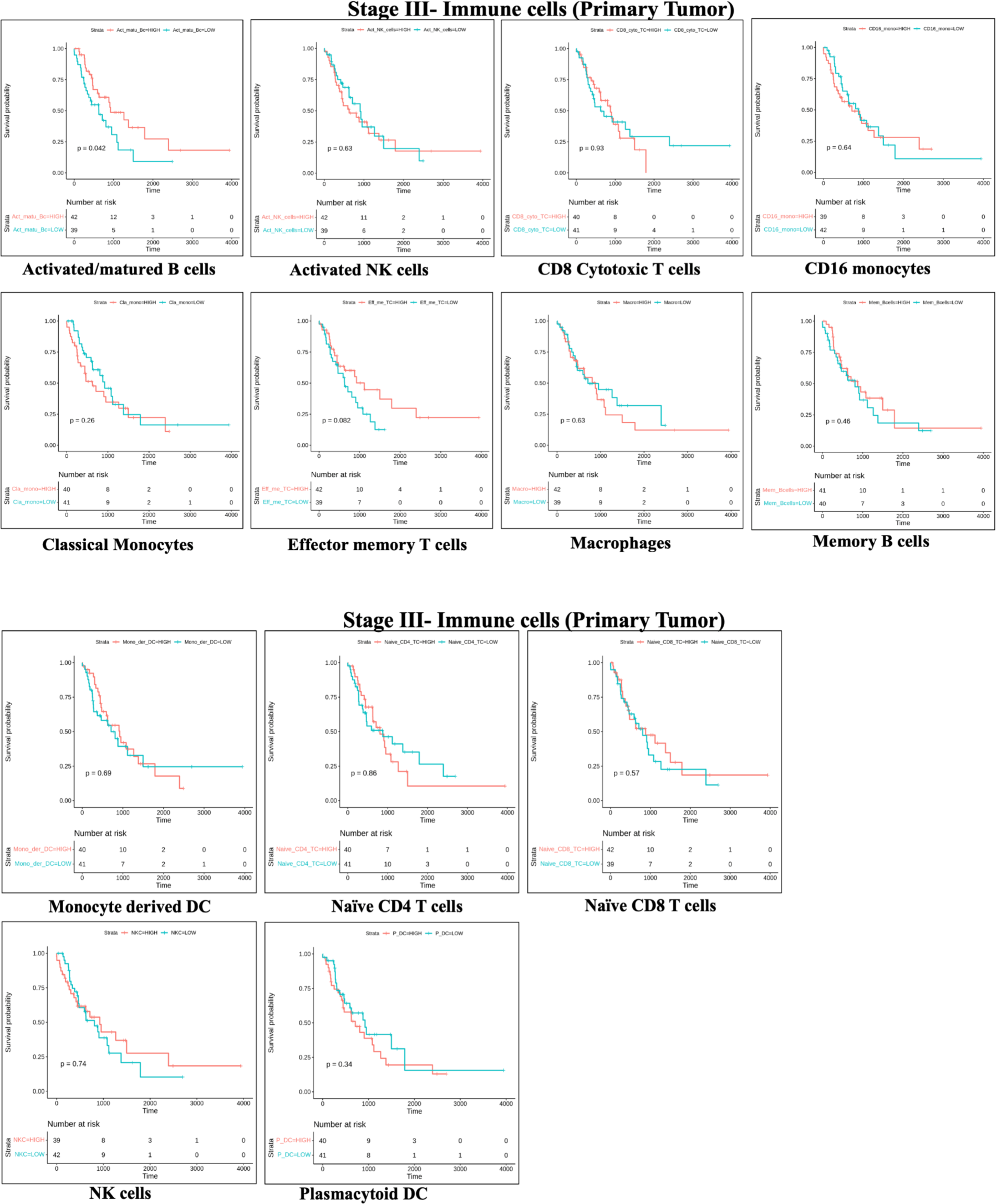
KM plot for immune cells in stage III of primary tumor.

**Figure 0.14.**
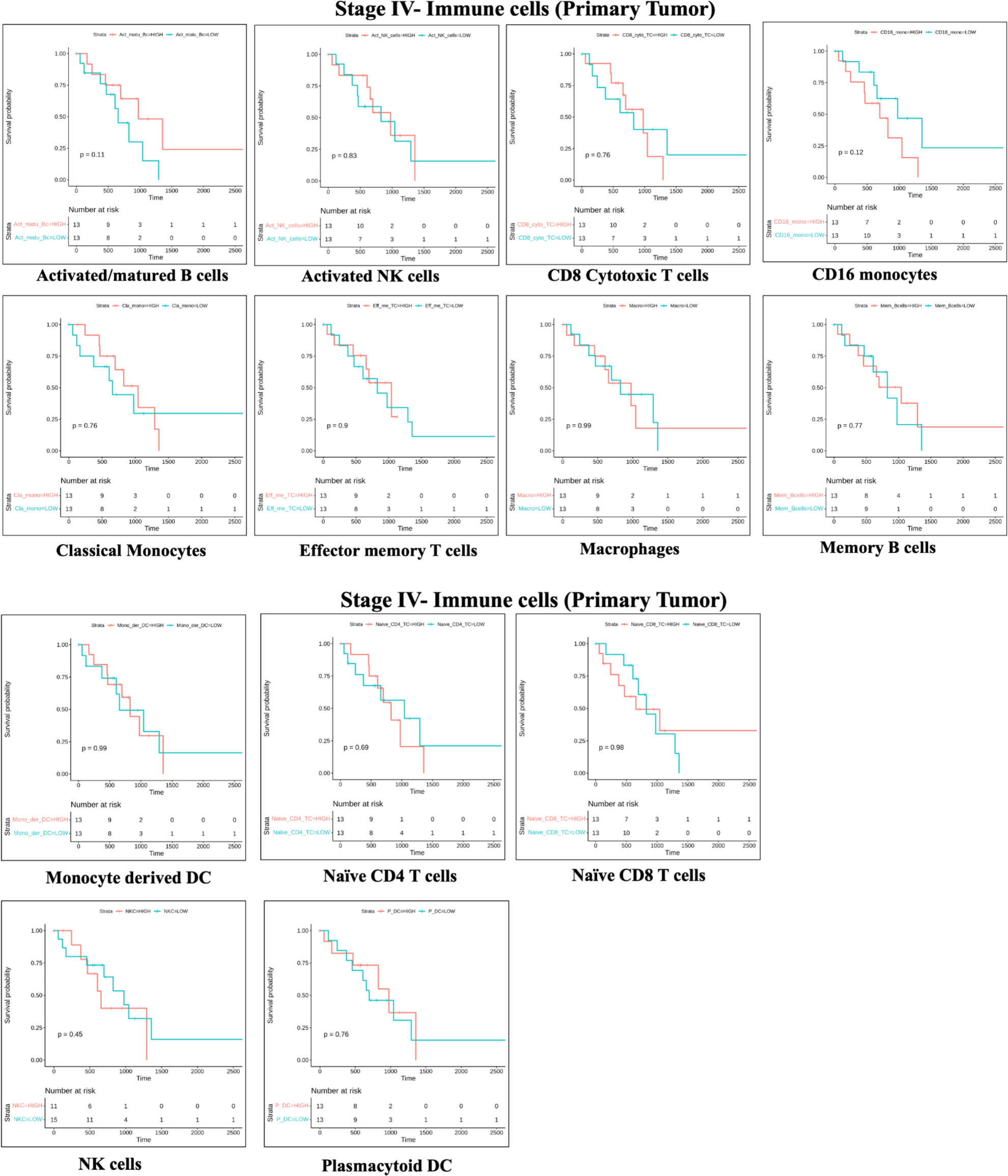
KM plot for immune cells in stage IV of primary tumor.

**Figure 0.15.**
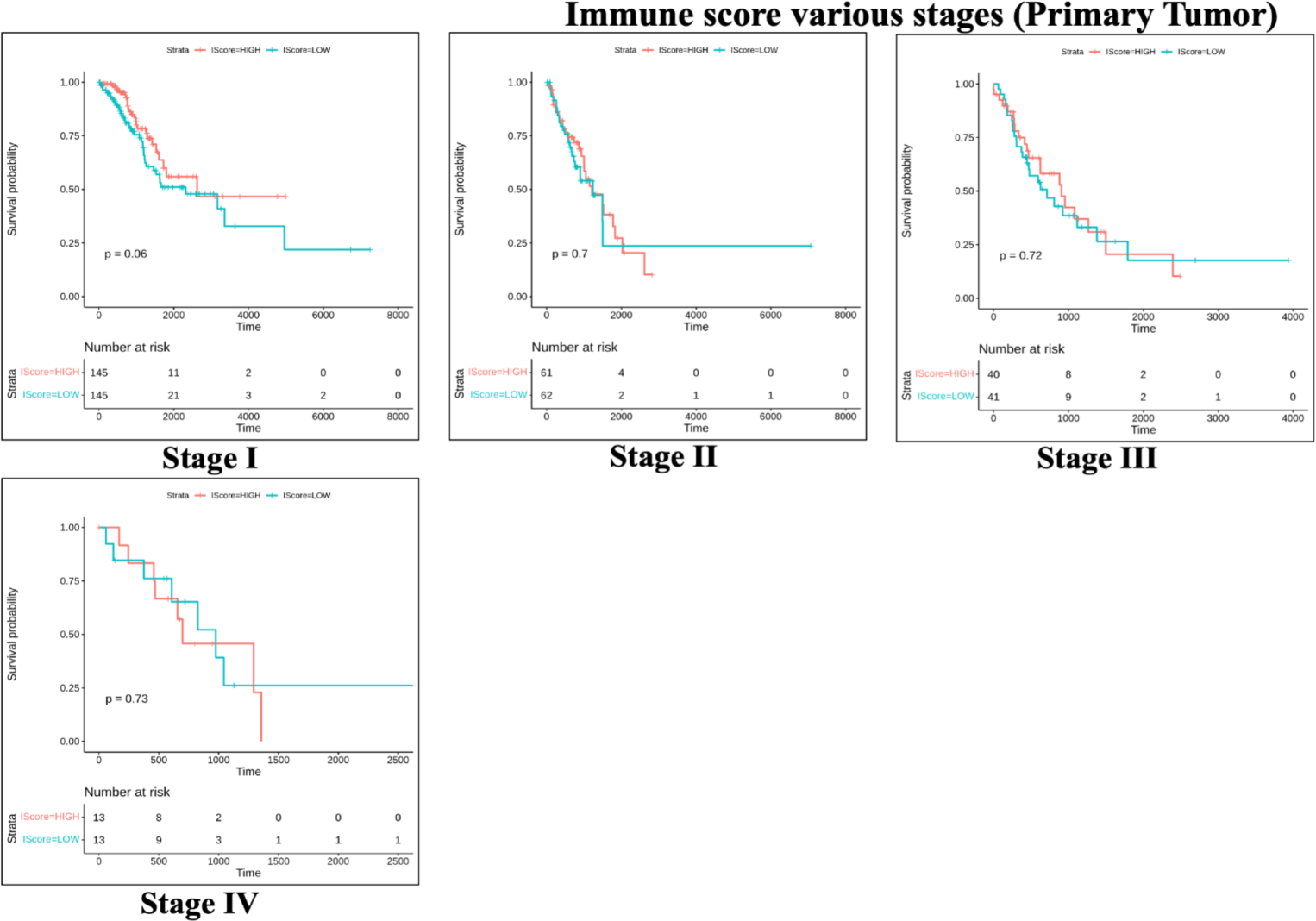
KM plot for immune score in different stages of primary tumor.

**Figure 0.16.**
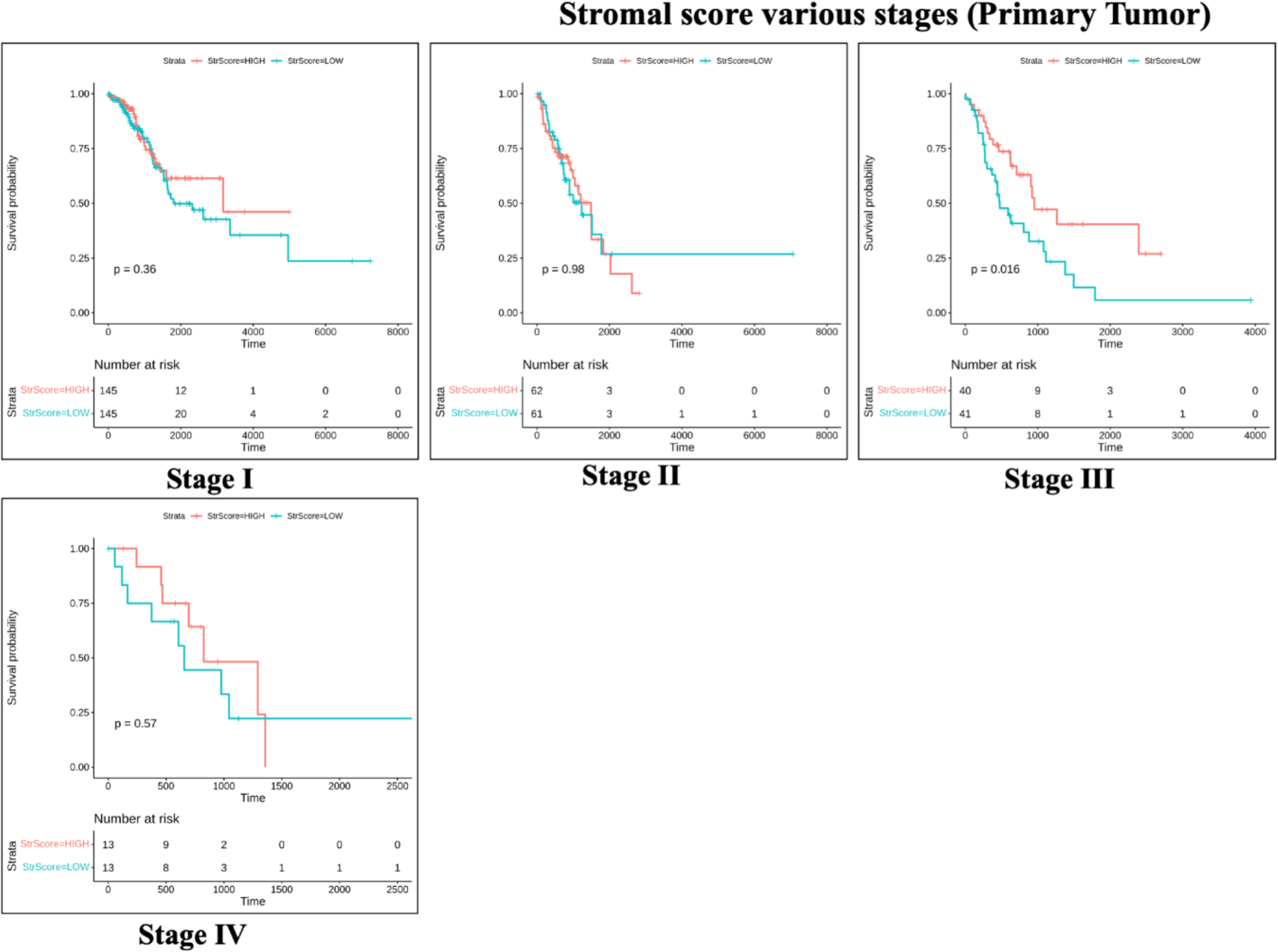
KM plot for stromal score in different stages of primary tumor.

**Figure 0.17.**
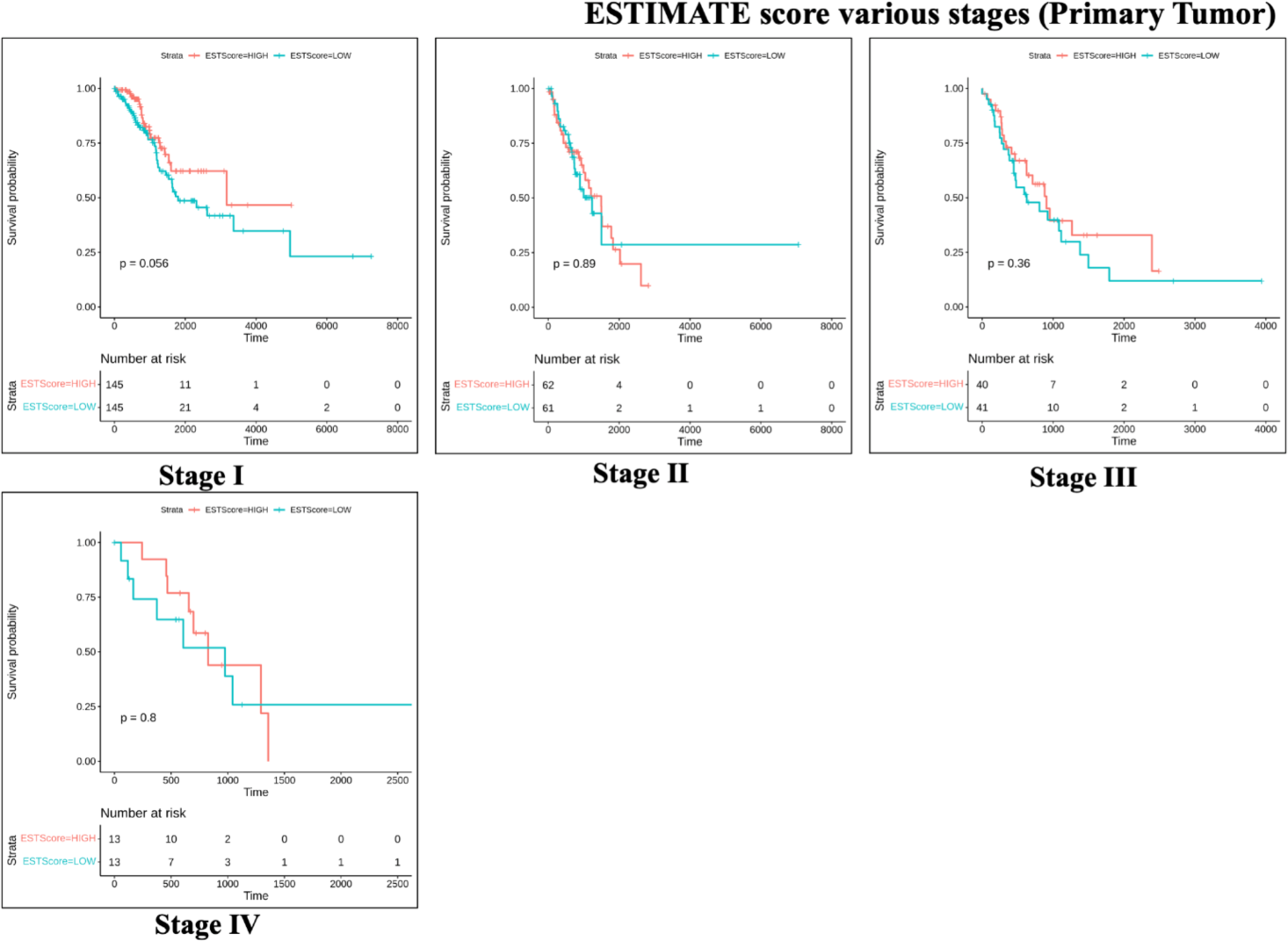
KM plot for ESTIMATE score in different stages of primary tumor.

## Abbreviations

AJCC: American Joint Committee on Cancer
BCR: B cell receptor
DCs: Dendritic cells
ESTIMATE: Estimation of Stromal and Immune Cells in Malignant Tumor Tissues Using Expression Data
ICD: O International Classification of Diseases for Oncology
KM: Kaplan-Meier
LC: Lung Cancer
LUAD: Lung Adenocarcinoma
NA: Not Available (non-significant in this case)
NK cells: Natural Killer cells
NOS: Not otherwise specified
NSCLC-non: small cell lung cancer
scRNA-seq: Single cell RNA sequencing
ssGSEA: Single-sample gene set enrichment analysis
T cells: T lymphocytes
T-regs: Regulatory T Cells
TCGA: The Cancer Genome Atlas
TIICs: tumor-infiltrating immune cells
TILs: Tumor infiltrating lymphocytes
TME: Tumor Microenvironment

## References

1. Borgan, R. (2001). Modeling Survival Data: Extending the Cox Model. Terry M. Therneau and Patricia M. Grambsch, Springer-Verlag, New York, 2000. No. of pages: xiii + 350. Price: $69.95. ISBN 0-387-98784-3. Statistics in Medicine, 20(13), 2053–2054. 10.1002/sim.956

2. Bremnes, R. M., Busund, L. T., Kilvær, T. L., Andersen, S., Richardsen, E., Paulsen, E. E., Hald, S., Khanehkenari, M. R., Cooper, W. A., Kao, S. C., & Dønnem, T. (2016, June). The Role of Tumor-Infiltrating Lymphocytes in Development, Progression, and Prognosis of Non– Small Cell Lung Cancer. Journal of Thoracic Oncology, 11(6), 789–800. 10.1016/j.jtho.2016.01.015

3. Colaprico, A., Silva, T. C., Olsen, C., Garofano, L., Cava, C., Garolini, D., Sabedot, T. S., Malta, T. M., Pagnotta, S. M., Castiglioni, I., Ceccarelli, M., Bontempi, G., & Noushmehr, H. (2016, May 5). TCGAbiolinks: an R/Bioconductor package for integrative analysis of TCGA data. OUP Academic. 10.1093/nar/gkv1507

4. Devarakonda, S., Morgensztern, D., & Govindan, R. (2015, July). Genomic alterations in lung adenocarcinoma. The Lancet Oncology, 16(7), e342–e351. 10.1016/s1470-2045(15)00077-7

5. Domingues, P., González-Tablas, M., Otero, L., Pascual, D., Miranda, D., Ruiz, L., Sousa, P., Ciudad, J., Gonçalves, J. M., Lopes, M. C., Orfao, A., & Tabernero, M. D. (2016, March). Tumor infiltrating immune cells in gliomas and meningiomas. Brain, Behavior, and Immunity, 53, 1–15. 10.1016/j.bbi.2015.07.019

6. Fridman, W. H., Pagès, F., Sautès-Fridman, C., & Galon, J. (2012, March 15). The immune contexture in human tumours: impact on clinical outcome. Nature Reviews Cancer, 12(4), 298–306. 10.1038/nrc3245

7. Ganesan, A. P., Johansson, M., Ruffell, B., Beltran, A., Lau, J., Jablons, D. M., & Coussens, L. M. (2013, August 15). Tumor-Infiltrating Regulatory T Cells Inhibit Endogenous Cytotoxic T Cell Responses to Lung Adenocarcinoma. The Journal of Immunology, 191(4), 2009–2017. 10.4049/jimmunol.1301317

8. Gentles, A. J., Newman, A. M., Liu, C. L., Bratman, S. V., Feng, W., Kim, D., Nair, V. S., Xu, Y., Khuong, A., Hoang, C. D., Diehn, M., West, R. B., Plevritis, S. K., & Alizadeh, A. A. (2015, July 20). The prognostic landscape of genes and infiltrating immune cells across human cancers. Nature Medicine, 21(8), 938–945. 10.1038/nm.3909

9. Herbst, R. S., Soria, J. C., Kowanetz, M., Fine, G. D., Hamid, O., Gordon, M. S., Sosman, J. A., McDermott, D. F., Powderly, J. D., Gettinger, S. N., K. Kohrt, H. E., Horn, L., Lawrence, D. P., Rost, S., Leabman, M., Xiao, Y., Mokatrin, A., Koeppen, H., Hegde P. S., … Hodi, F. S. (2014, November 26). Predictive correlates of response to the anti-PD-L1 antibody MPDL3280A in cancer patients - Nature. Nature. 10.1038/nature14011

10. Ji, R. R., Chasalow, S. D., Wang, L., Hamid, O., Schmidt, H., Cogswell, J., Alaparthy, S., Berman, D., Jure-Kunkel, M., Siemers, N. O., Jackson, J. R., & Shahabi, V. (2011, December 7). An immune-active tumor microenvironment favors clinical response to ipilimumab - Cancer Immunology, Immunotherapy. SpringerLink. 10.1007/s00262-011-1172-6

11. Kassambara, Kosinski, Biecek, & Fabian. (2021, March 9). Drawing Survival Curves using ggplot2. Drawing Survival Curves Using Ggplot2 • Survminer. Retrieved January 25, 2023, from https://rpkgs.datanovia.com/survminer/index.html

12. Ohue, Y., Kurose, K., Nozawa, R., Isobe, M., Nishio, Y., Tanaka, T., Doki, Y., Hori, T., Fukuoka, J., Oka, M., & Nakayama, E. (2016, November 30). Survival of Lung Adenocarcinoma Patients Predicted from Expression of PD-L1, Galectin-9, and XAGE1 (GAGED2a) on Tumor Cells and Tumor-Infiltrating T Cells. Cancer Immunology Research, 4(12), 1049–1060. 10.1158/2326-6066.cir-15-0266

13. Steen, C. B., Liu, C. L., Alizadeh, A. A., & Newman, A. M. (2020, January 21). Profiling Cell Type Abundance and Expression in Bulk Tissues with CIBERSORTx. PubMed Central (PMC). Retrieved January 25, 2023, from https://www.ncbi.nlm.nih.gov/pmc/articles/PMC7695353/

14. <αυτηορσ> The Cancer Genome Atlas Program. (n.d.). National Cancer Institute. Retrieved January 25, 2023, from https://www.cancer.gov/about-nci/organization/ccg/research/structural-genomics/tcga

15. Therneau, T. M. (2022, August 9). CRAN - Package survival. CRAN - Package Survival. Retrieved January 25, 2023, from https://cran.r-project.org/web/packages/survival/index.html

16. Tumeh, P. C., Harview, C. L., Yearley, J. H., Shintaku, I. P., Taylor, E. J. M., Robert, L., Chmielowski, B., Spasic, M., Henry, G., Ciobanu, V., West, A. N., Carmona, M., Kivork, C., Seja, E., Cherry, G., Gutierrez, A. J., Grogan, T. R., Mateus, C., Tomasic, G., … Ribas, A. (2014, November 26). PD-1 blockade induces responses by inhibiting adaptive immune resistance. Nature, 515(7528), 568–571. 10.1038/nature13954

17. Verma, M. (2024a). Assessing immune microenvironment in TCGA-LUAD via CIBERSORTx using single-cell derived signature matrix and ESTIMATE algorithm. bioRxiv, 2024.05.08.592760. 10.1101/2024.05.08.592760

18. Verma, M. (2024b). Unravelling immune cell signatures: CIBERSORTx-assisted construction of signature matrix from single-cell data. In bioRxiv. 10.1101/2024.05.05.592045

19. Yoshihara, K., Shahmoradgoli, M., Martínez, E., Vegesna, R., Kim, H., Torres-Garcia, W., Treviño, V., Shen, H., Laird, P. W., Levine, D. A., Carter, S. L., Getz, G., Stemke-Hale, K., Mills, G. B., & Verhaak, R. G. (2013, October 11). Inferring tumour purity and stromal and immune cell admixture from expression data. Nature Communications, 4(1). 10.1038/ncomms3612

20. Zheng, X., Hu, Y., & Yao, C. (2017). The paradoxical role of tumor-infiltrating immune cells in lung cancer. Intractable & Rare Diseases Research, 6(4), 234–241. 10.5582/irdr.2017.01059

